# Multimodal Learning of Pheromone Locations

**DOI:** 10.1101/2019.12.22.886564

**Authors:** Meenakshi Pardasani, Shruti D. Marathe, Maitreyee Mandar Purnapatre, Urvashi Dalvi, Nixon M. Abraham

## Abstract

Memorizing pheromonal locations is critical for many mammalian species as it involves finding mates and avoiding competitors. In rodents, pheromonal information is perceived by the main and accessory olfactory systems. However, the role of somatosensation in context dependent learning and memorizing of pheromone locations remains unexplored. We addressed this problem by training female mice on a multimodal task to locate pheromones by sampling volatiles emanating from male urine through the orifices of varying dimensions or shapes that are sensed by their vibrissae. In this novel pheromone location assay, female mice’ preference towards male urine scent decayed over time when they were permitted to explore pheromones v/s neutral stimuli, water. On training them for the associations involving olfactory and whisker systems, it was established that they were able to memorize the location of opposite sex pheromones, when tested 15 days later. This memory was not formed either when the somatosensory inputs through whisker pad were blocked or when the pheromonal cues were replaced with that of same sex. The association between olfactory and somatosensory systems were further confirmed by the enhanced expression of the Activity regulated cytoskeleton protein. Further, the activation of main olfactory bulb circuitry by pheromone volatiles did not cause any modulation in learning and memorizing non-pheromonal volatiles. Our study thus provides the evidence for associations formed between different sensory modalities facilitating the long-term memory formation relevant to social and reproductive behaviors.

## Introduction

In nature, when terrestrial rodents scour and encounter pheromones streaked on stones of different sizes and shapes, burrows and fallen leaves, how do they use different modalities to form a memory of the location of pheromones? Rodents are primarily macrosmatic mammals whose daily vital activities of foraging, finding mates and avoiding predators are dependent on olfaction (1). However, when it comes to memorizing scent marks, is their sense of smell enough or are associations being formed between multiple senses, integrating information to retain the memory of the marks? In rodents, coordinated action of sniffing and whisking has been observed during the exploratory behaviors (2, 3). However, it has not been tested if such multi-sensory actions facilitate the long-term memory formation of pheromones’ locations. We decided to undertake an approach for probing the role of different systems governing this by using a newly designed ‘multimodal pheromonal learning’ behavioral paradigm. Our paradigm lets the female mice use sensory information processed by whisker pathway to associate a location presented with volatile and non-volatile pheromones over a location presented with water, a neutral stimulus. Although the female mouse is allowed to have a direct exposure to soiled bedding, it can sample the volatiles emanating from male urine through a plate guarded by orifices of circular shapes with specific diameters or of triangular shapes. This paradigm offers us to probe for the role of whiskers and microvibrissae present on the snout along with the olfactory subsystems in forming the memory of pheromones location.

Classical knowledge reveals that pheromonal detection happens via vomeronasal organ (VNO) and main olfactory epithelium (MOE) which further project to accessory olfactory bulb (AOB) and main olfactory bulb (MOB), respectively. The output neurons project to bed nucleus of stria terminalis (BNST) and ‘vomeronasal amygdala’ and further to hypothalamic regions, which control the lordosis behavior in females (4–6). Using genetic and lesion-based approaches, it has been found that AOB senses non-volatile pheromones present in male urine and soiled bedding through its VNO receptors (Vomeronasal type 1, Vomeronasal type 2 and Formyl Peptide receptors) (7) while the trace amine-associated receptors (TAARs) present on MOE projecting to MOB mitral cells detect air-borne, volatile pheromones (8). Another receptor type includes MS4A receptors of “necklace” glomeruli subsystem found in caudal MOB, which are activated by certain pheromones such as 2,5-dimethylpyrazines (9). Such parallel mechanisms occurring via different olfactory subsystems reflect the complexity of pheromonal information processing by activating combination of receptor subtypes in rodents (10).

Male urine scent-marking is an excellent example of intra and inter-sex social communication (11) leading to display of dominance by the giver (12, 13) or attractive/preferential behaviors by recipient females (14). In the context of mating, pheromones are important to be learnt and remembered by female mice. Earlier studies suggested that repeated vagino-cervical stimulation along with the pheromonal exposure is required for acquisition of long-term memory in females (15). Although, now we know that a single exposure is enough to form a memory (14), we do not yet understand the involvement of different sensory modalities in governing the memory formation/acquisition.

Our paradigm involves probing the effect of pheromone priming on the chemo-as well as whisker-mediated investigatory behavior and long-term memory formation in female mice. We observed that the preference towards a zone containing pheromones decayed over time when mice were freely exploring the arena during the initial 4 days. Further, we trained female mice to learn the associations involving male pheromones, both volatiles and non-volatiles, and whisker-mediated sensory cues. These mice exhibited long-term memory when tested 15 days after the training, in the absence of any sensory cues. In order to investigate if somatosensation is important for locating such marks, we checked if the preference towards male mouse pheromones is negatively affected when the sensory input through whiskers and skin is blocked during the 15 days of training. Interestingly, we saw poor pheromonal long-term memory in these females suggesting the involvement of somatosensory system along with main and accessory olfactory systems. To investigate the differential activity-dependent activation across associative brain regions under sensory intact and deprived conditions, we examined the expression of Activity regulated cytoskeleton (Arc) protein. Significantly higher Arc protein expression was found in the somatosensory cortices and hippocampi of mice whose sensory inputs were not blocked. Such an increased and specific activation of brain regions support our finding of the robust pheromonal memory formation in the whisker-intact group of mice. To probe if pheromone-dependent activation of main olfactory system can modulate other olfactory-driven behaviors, we carried out volatile discrimination learning and memory tasks. The pheromonal exposure induced Whitten effect, however, did not cause any differences in the MOB mediated discrimination learning pace for various non-pheromonal volatiles. Our data thus provide evidences for novel associations between different sensory modalities in facilitating long-term memory formation of pheromones.

## Methods

### Subjects

A total of 76 C57BL/6J female (10-14 weeks, Jackson Laboratories) and 8 C57BL/6J male mice (12-14 weeks, Jackson Laboratories) were utilized for the study. Three different groups of female mice were tested on ‘multimodal pheromonal learning’ paradigm. Group 1 consisted of eight whisker intact female mice trained for associating Male soiled Bedding and Urine volatiles (MBU) with a particular hole diameter punched on the plate guarding the chamber 1, where the stimulus is kept. Group 2 had eight female mice, whose whiskers were trimmed during the training phase for the associations involving MBU. But with the same group, whiskers were kept intact during initial testing and memory testing phases. Group 3 consisted of eight whisker intact female mice trained for associating Female soiled Bedding and Urine (FBU) with a particular hole diameter. Group 4 consisted of eight whisker intact female mice trained for associating MBU and neutral stimuli with the distinct shape (either triangle or circle of perimeter/circumference of 31.4mm) of the plate guarding the chamber. For Arc immunohistochemical analyses, a total of fourteen mice encompassing the whisker intact and deprived groups were trained. Eight of these mice were used to investigate Arc expression on the memory day 15^th^. All the abovementioned groups were trained using the same paradigm. Four mice each from group 1,2 and 3 were utilized for measuring sniffing behavior towards pheromonal volatiles. For a given experiment, male urine was collected from either one male mouse or two mice housed together (C57BL6/J, aged 10-12 weeks), who never had a sexual experience. Female urine was collected from two female mice housed together (C57BL6/J, aged 10-12 weeks), who were also never sexually experienced. Soiled bedding was collected from the respective cages, just before the commencement of the experiment.

For non-pheromonal odor discrimination training, 30 female mice, aged 8-14 weeks, were used. These were divided equally in two groups, one group (experimental, Group 5) exposed to MBU every-day during the discrimination training period while the other group (control, Group 6) unexposed. 12-hours light/dark cycle was maintained and mice were grouped in individually ventilated cages in a temperature- and humidity-controlled animal facility. All animal care and procedures were in accordance with the Institutional Animal Ethics Committee (IAEC) at IISER Pune and the Committee for the Purpose of Control and Supervision of Experiments on Animals (CPCSEA), Government of India.

#### Multimodal Pheromonal learning

##### Apparatus

It consisted of an arena (60cm x 30cm x 15cm, length x width x height, coated with non-reflective black spray paint), which was divided into 3 equally spaced zones. Two 10cm x 10cm x 15cm chambers guarded by removable plates were placed at the opposite ends of the length of arena. One removable plate guarding a particular chamber had equidistant 10mm diameter orifices on it, keeping other three sides of each chamber devoid of any orifices. Chamber on the opposite side was guarded by a removable plate having equidistant 5mm diameter orifices. This allows the animal to sample the volatiles emanating from the chamber through only one (front) side. One of each kind guards these chambers when the animal was sampling the volatiles during the entire experimental duration (initial testing, training and memory testing). The apparatus was custom-built using black acrylic sheets (6mm thickness for the walls and 10mm thickness for the base). Each chamber contained a 55mm petri-dish filled with 100μL urine (attractive pheromonal stimulus in chamber 1) or water (neutral stimulus in chamber 2) (Figure 1B). Two different apparatuses were used for carrying out the multimodal pheromonal learning paradigm in the MBU and FBU groups. For shape-based multimodal pheromonal learning task, one of the removable plate guarding one chamber had triangle shaped orifice while the other plate had circle shaped orifices of 10mm diameter. The orifice dimensions were chosen such that the circumference of the circular orifices matches with the perimeter of the triangular orifice on the opposite sides.

**Figure 1.**
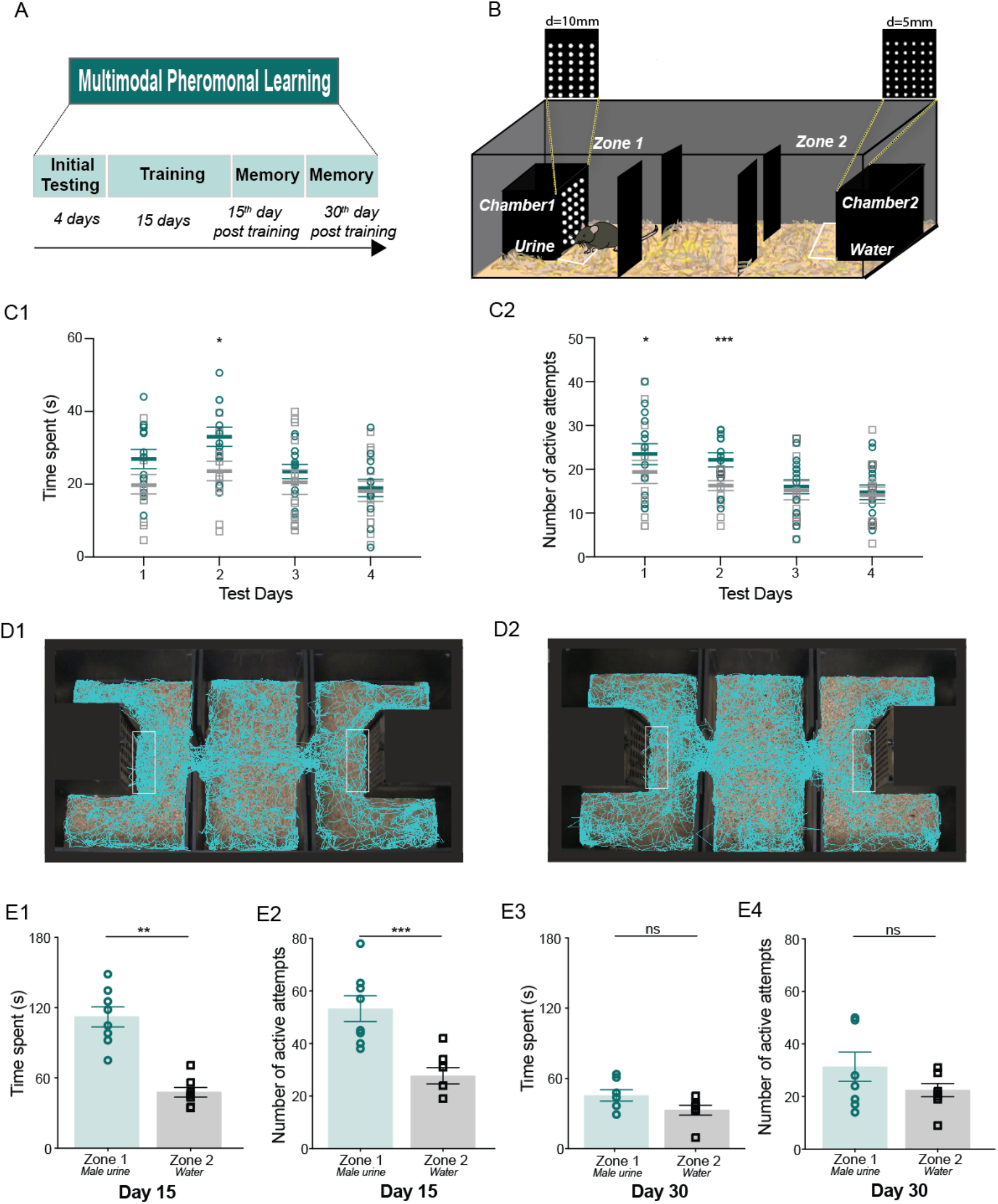
Multimodal associative learning causes long-term memory formation of pheromone locations in female mice. A. Experimental design of ‘multimodal pheromonal learning’ paradigm. It involves an initial testing phase of 4 days, training phase of 15 days and the testing of memory at 15^th^ and 30^th^ day post training. B. Diagrammatic representation of the set-up used for the assay. Female mouse is shown to be sensing the volatiles emanating through the specific diameter orifices from the stimulus present in OSP chamber. Perimeter demarcated by white line represents the zone where time spent by mouse is calculated using the EthoVision software by tracking the ‘nose-point’ of the animal. C1. Similar time spent near OSP chamber (teal colored symbols) v/s NS chamber (gray colored symbols) on day 1, 3 and 4 of testing phase. Increased time spent on day 2 indicates emerging preference towards OSP, which, declines as the days of exploration progress. (p > 0.05, F = 4.34 Repeated Measures (RM) Two-way ANOVA; Bonferroni’s multiple comparison test, p < 0.05 for day 2, p = 0.1 for day 1, p > 0.9 for day 3 and day 4; N = 13 mice). C2. Similar number of active attempts on plate guarding OSP chamber v/s NS chamber on day 3 and 4 during initial testing phase. On day 1 and day 2, mice exhibited more number of active attempts manifesting initial preference towards OSP, which was reduced on day 3 and 4 (p > 0.05, F = 4.1 RM Two-way ANOVA; Bonferroni’s multiple comparison test, p = 0.016 for day1, p = 0.0003 for day 2, p > 0.9 for day 3 and day 4; N = 15 mice) D1, D2. Tracks recorded using EthoVision software by tracking the ‘nose point’ of the animal. Visibly more time spent near zone 1 compared to zone 2 in case of memory day 15^th^ (D1) while equal time spent for both the zones in case of memory day 30^th^ (D2) is seen. The area demarcated is used for calculating time spent near the two chambers. E1. More time was spent near OSP chamber than NS chamber on day 15 post training period (p = 0.007, Wilcoxon matched pairs signed rank test, N = 8 mice). Memory was tested in the absence of any stimuli (urine, soiled bedding and water). E2. Number of active attempts on plate guarding OSP chamber was higher than on NS chamber on day 15 post training period (p = 0.0004, Paired two-tailed student’s t-test, N = 8 mice). Memory was tested in the absence of any stimuli (urine, soiled bedding and water). E3. Similar amount of time was spent near OSP chamber and NS chamber on day 30 post training period (p = 0.109, Wilcoxon matched pairs signed rank test, N = 7 mice). Memory was tested in the absence of any stimuli (urine, soiled bedding and water). E4. Similar number of active attempts were made on plate guarding OSP chamber and NS chamber on day 30 post training period (p > 0.1, Wilcoxon matched pairs signed rank test, N = 8 mice). Memory was tested in the absence of any stimuli (urine, soiled bedding and water).

##### Paradigm

The experimental design involves an initial testing phase of 4 days, training phase of 15 days and the testing of memory at 15^th^ day and 30^th^ day post training (Figure 1A). The initial testing phase was to investigate if female mice exhibit innate preference towards a zone presented with attractive volatile and non-volatile pheromones from male mice (zone 1) over a zone containing water stimulus (zone 2). To remove any directional bias towards a particular zone, the apparatus was rotated by 180° everyday during the initial testing and training phases. Mice were counterbalanced for the association with a specific hole diameter for circular shapes and the volatile cue pairing (Urine/5mm v/s Water/10mm and Urine/10mm v/s Water/5mm, Groups 1, 2 and 3) and for triangular or circular shapes and volatile cues (Urine/triangular shaped orifices v/s Water/circular shaped orifices and Urine/circular shaped orifices v/s Water/triangular shaped orifices, Groups 4) to remove any possible bias towards specific hole diameter or shapes. The chambers remained the same during this counter-balancing, although new plates were used. Training was done following the initial testing phase for a period of 15 days. Each day, the animal was restricted in both the zones for a period of 15 minutes each (alternating between the two zones after every 5 minutes). To check for the multimodal memory on 15^th^ and 30^th^ day, all volatile and non-volatile pheromonal stimuli (urine and the soiled bedding with non-volatile pheromonal traces) and neutral water stimuli were removed from the chambers of the apparatus while keeping the plates of specific diameter orifices undisturbed. The time spent in each of the zones, specifically in front of chamber 1 and chamber 2 were calculated using EthoVision software (Noldus Information Technology). The nose point feature (used to track animal) on EthoVision is used to visualize the tracks taken by an animal (Figure 1 D1 and D2) and to calculate the time spent. Number of active attempts on the plates guarding chamber 1 and 2 were calculated manually by considering single nose poke into the hole as one attempt.

In case of testing the involvement of somatosensation in pheromone location learning and memory, an anesthetic gel (Lignocaine Hydrochloride gel-Lox-2% jelly, Neon laboratories) was applied on the snout of the group 2 mice during the training phase of 15 days. Whisker trimming was done at a frequency of five days from the start of the training phase. Whiskers were allowed to grow and were never trimmed again after the training period. Mice had re-growing whiskers when the day 15 memory was tested, thereby allowing us to specifically check for the role of somatosensation in mediating the association of pheromonal cue with hole-size.

#### Go/no-go odor discrimination

##### Odors

Odors used were 1,4-Cineol (CI), Eugenol (EU), Amyl acetate (AA), Ethyl butyrate (EB), Benzaldehyde (BZ), Nonanol (NN), Hexanal (HX) and 2-Pentanone (PN). The odors were diluted to 1% in mineral oil and further diluted 1:20 by airflow. All odors were bought from Sigma-Aldrich and mineral oil was bought from Oswal Pharmaceuticals, Pune, Maharashtra, India.

##### Apparatus

All experiments were done using two custom-made eight channel olfactometers controlled by software written in Igor Pro (Wavemetrics, OR). Briefly, mouse was put into an operant chamber, which had a combined odor sampling and reward delivery port on one end of the chamber. This ensured tight association of reward with the odor presented in a trial. Inside the odor port, a lick tube was placed on which the animal needs to lick to get the reward. Insertion of animal’s head in the port resulted in breaking of an IR beam guarding odor/reward port, which initiated the trial. The odor and final (diversion) valves controlled the flow of odor in a time-dependent manner, allowing opening of final valve 500ms after odor valve opening. Each rewarded (S+) and non-rewarded (S-) odor was presented through either of the two valves. A total of four odor valves for two odors were used. The apparatus and the task design followed is same as done in previous published studies (Figure 3C1, C2, C3) (16–19)

**Figure 2.**
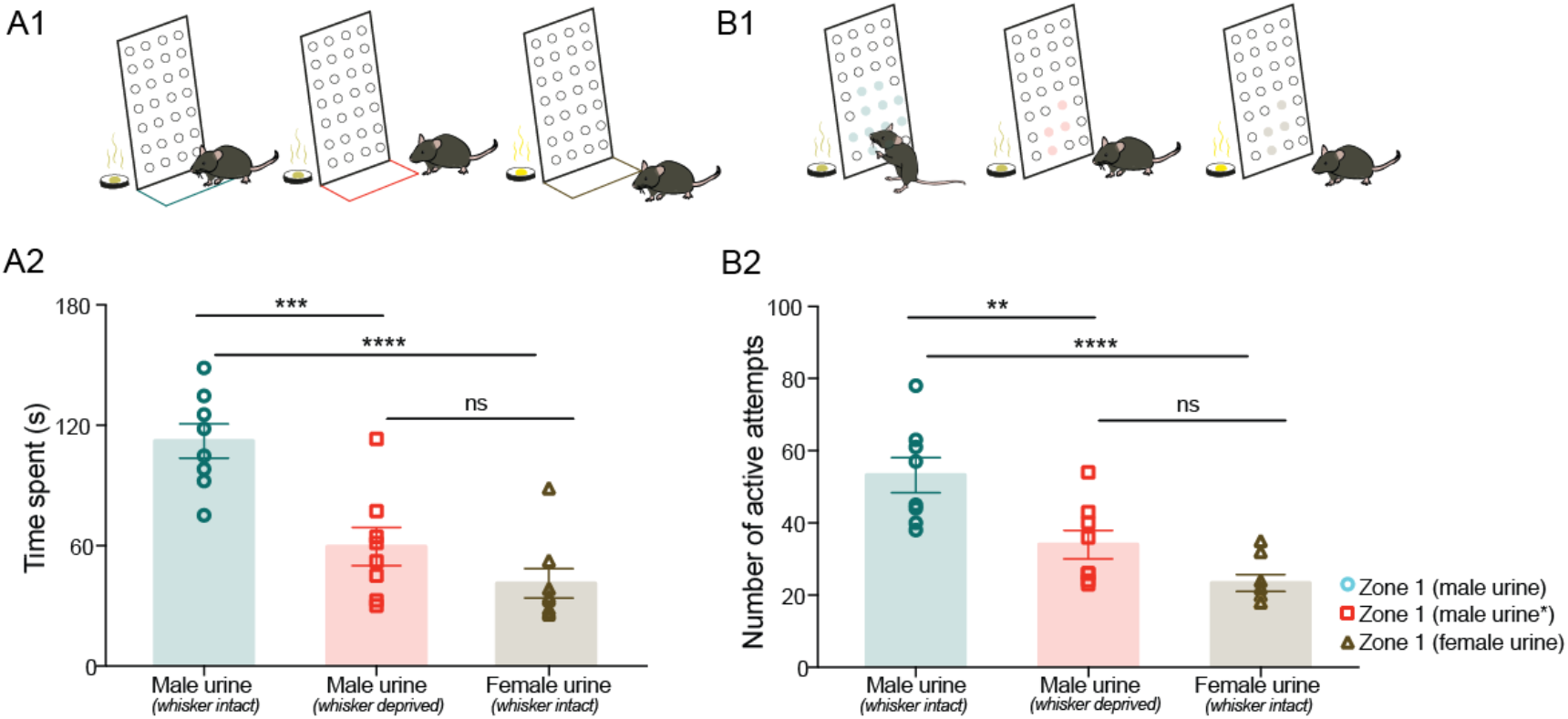
Long-term memory of pheromone location is facilitated by the association formed between olfactory and whisker systems. A1. Schematic representing a female mouse sampling in front of the plate guarding OSP chamber in the perimeter demarcated by a colored line. Nose point of the mouse snout is tracked for measuring parameter “time spent” by EthoVision software (Teal: whisker intact mouse trained towards male MBU, Red: whisker deprived mouse towards MBU and Brown: whisker intact mouse towards FBU). Memory was tested in the absence of any stimuli (urine, soiled bedding and water). A2. Reduced memory was observed for whisker deprived mice trained with male urine and whisker intact mice trained with female urine compared to whisker intact mice trained with male urine on 15^th^ day memory testing (p < 0.0001, F = 18.69, Ordinary One-way ANOVA, Bonferroni’s multiple comparison test; Male urine v/s Male urine*: p = 0.0008, Male urine v/s Female Urine: p < 0.0001, Male urine* v/s Female urine: p > 0.1, N = 8 mice for all groups) (* whisker deprived). B1. Schematic representing a female mouse making active attempts by poking its snout multiple times into the orifices of a particular diameter on plate guarding OSP chamber. Color of the orifices’ circumference depict the experimental condition; teal: whisker intact group trained towards male urine; red: whisker deprived group towards male urine and brown: whisker intact group towards female urine. Memory was tested in the absence of any stimuli (urine, soiled bedding and water). B2. Reduced number of active attempts were observed for whisker deprived mice trained with male urine and whisker intact mice trained with female urine compared to whisker intact mice trained with male urine on 15^th^ day memory testing (p < 0.0001, F = 15.27, Ordinary One-way ANOVA, Bonferroni’s multiple comparison test; Male urine v/s Male urine*: p < 0.01, Male urine v/s Female Urine: p < 0.0001, Male urine* v/s Female Urine: p > 0.1, N = 8 mice for all groups) (*whisker deprived).

**Figure 3.**
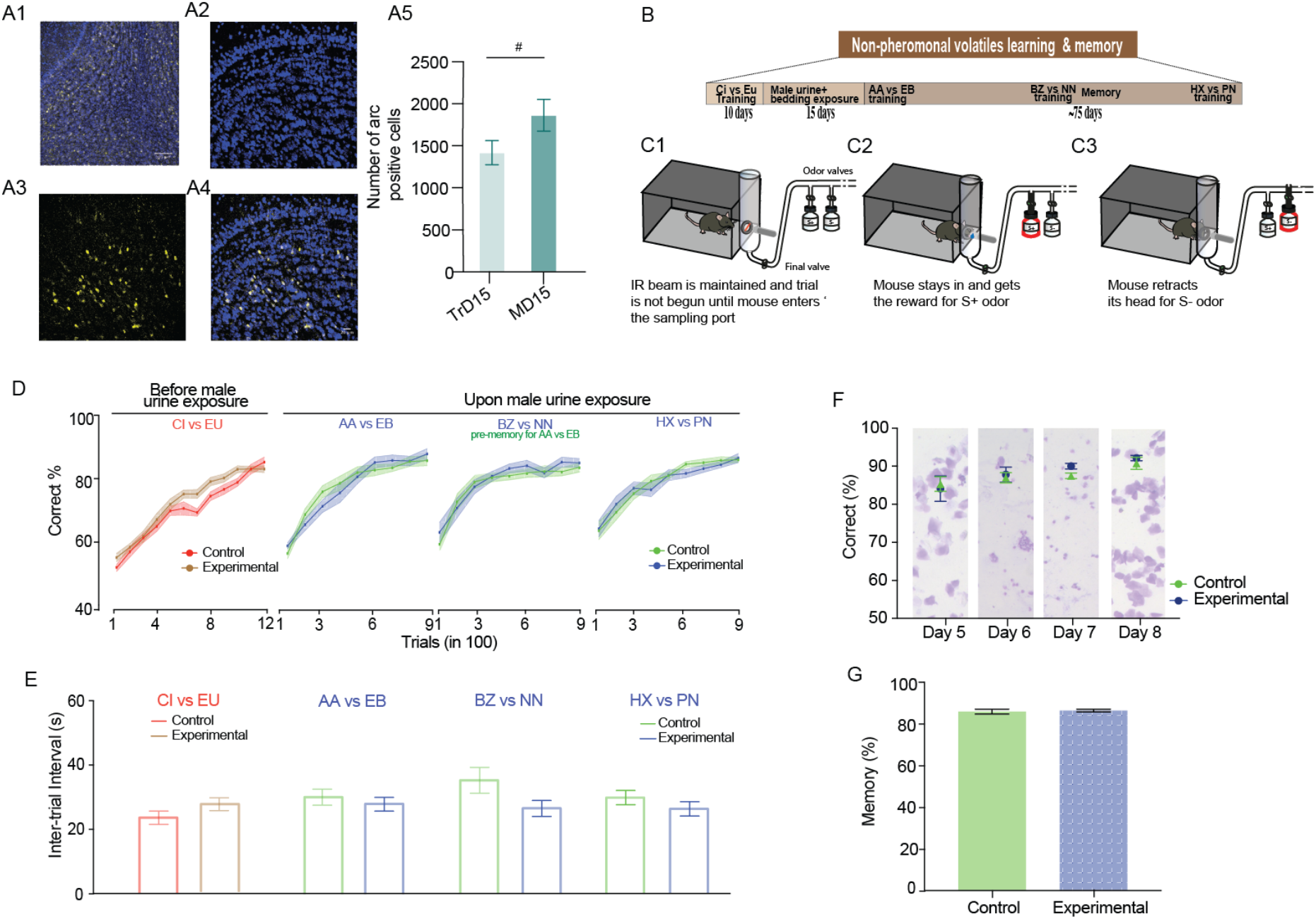
MOB circuitry activation does not modulate non-pheromonal volatile discrimination learning and memory. A1. Arc immunoreactivity in the MOB of whisker intact mouse exposed to MBU. Blue: DAPI, Yellow: Arc, Scale Bar: 100μm. A2, A3, A4. Magnified images of the granule cell layer of the MOB, Scale bar: 20μm. A5. A trend in increase in the proportion of Arc immunoreactive cells in the MOB of whisker intact group from TrD 15 to MD15 (#, p = 0.072, Unpaired t-test, twotailed). B. Experimental design for the go/no-go associative olfactory learning training for MBU exposure and no-exposure female mice groups to investigate their olfactory discrimination learning and memory. C1, C2, C3. Schematic of go/no-go odor discrimination paradigm indicating the sequence of events in a rewarded (A2) v/s unrewarded trial (A3) for a trained mouse. D. Learning curves depicting accuracy of discrimination across different odor pairs before MBU and after MBU exposure, shown as percent correct choices of 100 trials. Mice from both the groups were trained for CI v/s EU to ensure similar learning efficiencies before Whitten effect was induced (Ordinary Two-way ANOVA, Bonferroni’s multiple comparison test, p > 0.05 for all data points in learning curve). No difference in the pace of learning across different odor pairs was observed after Whitten effect was induced (AA v/s EB: p > 0.1, F = 0.12; BZ v/s NN: p > 0.1, F = 1.43; HX v/s PN: p > 0.1, F = 0.19; Ordinary Two-way ANOVA, Bonferroni’s multiple comparison test, p > 0.05 across all data points for all learning curves, N = 11-15 mice for both groups across odor pairs). E. Inter-trial interval for both groups across different odor pairs. It gives an indication of motivation of carrying out the trials, i.e. motivation is similar in both the groups. CI v/s EU: p = 0.2, Mann-Whitney test; AA v/s EB: p = 0.42, BZ v/s NN: p = 0.06, HX v/s PN: p = 0.26, Unpaired two-tailed student’s t-test, N = 11-15 mice for both groups across odor pairs). F. Average discrimination accuracy values during four consecutive days of AA v/s EB training done after the induction of Whitten effect. On each of these days, the accuracy was similar between experimental and control group. Vaginal cytology in background for each days on the plot depicts induction and synchronization of estrous cycle for experimental group (estrous: day 5, metestrous: day 6, diestrous: day 7 and proestrous: day 8) (p > 0.1, F = 0.49, Ordinary Two-way ANOVA; Bonferroni’s Multiple comparison test, p > 0.9 across all data points, N = 7 mice) G. Memory for AA v/s EB odor pair was checked one month after the odor pair learning was performed by mice. Memory was similar across both the groups. (p > 0.1, Mann-Whitney test, N = 13 for both groups)

##### Task habituation phase

First week post-water restriction involved training with the standard operant conditioning procedures. Initially, mice were rewarded with water upon insertion of head into the sampling port and breaking the IR beam. The difficulty levels increased gradually, wherein the animal needed to lick on the tube to get the reward. At the end of this phase, odor valve was introduced and the air passing through the mineral oil (solvent used for odorants) bottle was used as the stimulus.

##### Discrimination training phase

As the animal initiated the trial by breaking the beam, one of the odor valves and a diversion valve (DV) open for certain duration. The DV helps to divert the odorized air to the exhaust for 500ms and thereafter to the odor port for the pre-decided stimulus duration (2s for the odor discrimination training). For a rewarded (S+) odor to be correctly registered, animal needs to lick any 3 out of 4 bins of 500ms to get the reward. For an unrewarded odor (S-) to be correct, animal is allowed to lick at most 2 out of 4 bins. No punishment is given for a wrong response towards S-odor. There is a fixed inter-trial interval (ITI) of 5s during which the animal cannot initiate the next trial. Usually an optimally motivated animal took more than 10s as the ITI. Odors were presented in a pseudo-randomized manner (not more than 2 successive presentations of same odor and S+/S-odors were distributed equally in a block of 20 trials).

##### Resistance to memory extinction task

Once the animals reached a criterion performance of >80% for an odor pair for which long-term memory needs to be checked, we performed resistance to memory extinction task. This was done to stabilize the memory of an odor pair and is based on the ‘Partial reinforcement theory’ (20). The task comprised of 100 trials with 50 trials for S+ and S-each. The S+ trials are reinforced pseudo-randomly only for half of the trials as compared to a normal reinforcement learning where all S+ trials are rewarded. Such a partial reinforcement learning increases the attention of the animal as the factor of anticipating a reward is no longer true for some trials and that the trial outcomes cannot be expected by the animal. This increases the association strength and helps preventing memory extinction.

##### Memory task

The olfactory memory was checked for AA v/s EB, 30 days after carrying the resistance to memory extinction task. The memory task comprised of 200 trials, first 60 trials of background odor pair and subsequent 140 trials with interleaved memory trials [4 trials (two S+ and two S-trials) in every block of 20 trials of background odor pair]. The memory task was performed only after the accuracy of background odor pair reached >80% for all animals. For checking AA v/s EB memory, BZ v/s NN was used as a background odor pair. Memory trials were not rewarded. Memory is calculated as an average accuracy percent of 28 trials (14 S+ and 14 S-trials) tested over a total of 140 trials.

##### Sniffing behavior towards pheromones

To investigate the sniffing strategies of female mice towards male urine and female urine volatiles under whisker trimmed v/s intact conditions, a head-restraining method was used to precisely deliver the volatiles on the snout of the mice (21). These were subsets of mice utilized for multimodal pheromonal learning assay and their sniffing strategies were tested 2 months after their 30^th^ day memory was investigated. They were implanted with a head-post (custom-built head fixation set-up) on their head. The skin overlaying the skull was removed and head post was fixed using dental cement (Tetric N-Ceram, Ivoclar Vivadent). Surrounding exposed skull was covered using the dental acrylic cement (DPI RR Cold cure, acrylic repair material). This surgery was performed under anesthesia maintained by intraperitoneal injection of Ketamine (Celtamin, Celtiss therapeuticals) 60mg/kg and Xylazine (Vea Impex) 10mg/kg. The eyes were hydrated with 1% (w/v) carboxymethylcellulose (Refresh liquigel, Allergen India) to prevent dryness. Recovery time of two days was given and the weights were measured every day. Upon recovery, they were placed in a plastic tube and headpost was screwed onto a metallic device fitted on a platform. Urine volatiles and air presentations were done in a pseudorandom fashion through a nozzle. 100μL of urine was used for the same. Their sniffing behavior towards the volatiles was recorded using airflow pressure sensor (AWM2300V, Honeywell) placed near one of the nostrils (21). Each presentation was carried out for a period of 2s repeated over 10 such presentations of the same stimulus (total 20 presentations), accounting for a total of 20s time of exposure to pheromonal volatiles on a single day.

##### Immunohistochemistry

Mice were sacrificed using Thiopentone (Thiosol sodium, Neon laboratories, 50mg/kg) and perfused using a fixative. Transcardial perfusion was done using 50mL of 1X Phosphate buffered saline (PBS) followed by 4% (w/v) Paraformaldehyde (PFA). Brain of the animal was dissected and incubated in 4% PFA overnight at 4^0^C. Next day, tissue was washed twice with 1X PBS to remove extra fixative. In order to carry out cryotome sectioning, brain was cryopreserved using 20% sucrose (w/v) for a day and put on a rotator (Tarsons rotaspin). It was then transferred to 30% sucrose (w/v) at 4^0^C overnight. Brain tissue was embedded in optimal cutting temperature (OCT) medium (Leica,14020108926) and sectioned in a cryotome. 50μ m coronal sections were cut. Sections were washed in 1X Tris Buffered Saline (TBS) twice for 10 minutes each in a 6-well plate. Blocking solution (5% Bovine Serum Albumin (BSA) and 1% Triton-X in TBS) was added to the wells and plate was kept undisturbed for 1.5 hours. Sections were then incubated in primary antibody (Rabbit anti-Arc, Arc-156003, Synaptic Systems) diluted 1:1250 in blocking solution (1% BSA, 0.1% Triton-X in TBS) for 14 hours in 4^0^C. TBS washes (thrice, 15 minutes each) were given to remove the excess of antibody. Incubation with secondary antibody (Anti-Rabbit Alexa Fluor 594, Jackson’s immunoresearch) diluted 1:500 in 1% BSA in TBS and kept for 2 hours at room temperature. Sections were then washed with TBS (thrice, 15 minutes each) and labelled with DAPI (Sigma, 1:500). They were mounted on glass slides using Vectashield anti-fade mounting medium (Vector labs, H-1000).

##### Confocal Imaging

For quantification of Arc labelled nuclei across the whole brain coronal sections, imaging using SP8 confocal microscope (Leica) was done. Quantification of the Arc-positive cells was done using Imaris software (Oxford Instruments).

##### Statistical analyses

All data are represented as Mean ± SEM (Standard Error of Mean) and analysed using Graphpad Prism 8.0 (Graphpad Software Inc, USA). Normality of the data was checked using Shapiro-Wilk test. For data that followed normal distribution, Unpaired student’s t-test, Paired student’s t-test, One-way Analysis of Variance (ANOVA) and two-way ANOVA with *post hoc* Bonferroni’s multiple comparison testing were carried out wherever required. All t-tests were twotailed. For data that did not follow normal distribution, Mann-Whitney and Wilcoxon matched pairs signed rank tests were carried out wherever required. * represent p < 0.05, ** p < 0.01, *** p < 0.001 and **** p < 0.0001.

## Results & Discussion

In their natural habitats, rodents may locate scent marks associating with different surrounding objects of various sizes, shapes and textures which raises the possibility for the stimulation of whiskers and microvibrissae during this process. Can then, the memory formation for location of scent mark be facilitated by the activation of whisker system along with the olfactory subsystems? In order to answer this, we designed a novel ‘multimodal pheromonal learning’ paradigm to test the preference of female mouse towards the urine of opposite sex (Figure 1A). It involved a set-up consisting of zone 1 and zone 2. A 55mm petri dish containing male urine was kept in the chamber 1 [used henceforth as opposite sex pheromones (OSP) chamber] of zone 1. Neutral stimulus (NS), water was kept in a petri dish in chamber 2 [used henceforth as NS chamber] present in zone 2. A mouse can sample the volatile stimuli through specific diameter orifices made on one side of the chamber as depicted in figure 1B. To avoid any bias towards specific chamber, hole diameters were counterbalanced across mice for sensing male urine and water. Separate plates of a particular hole diameter were used in these conditions.

To investigate mice’ (Group 1) innate preference towards any zones, they were allowed to explore both zones for a total of 10 minutes everyday during the initial 4 testing days. To remove any possible directional bias shown by the mice towards a particular zone, the apparatus was rotated by 180° everyday during the initial testing and training days (See methods). The preference was quantified based on the time animal spent in front of OSP chamber sensing the non-volatile pheromones from male soiled bedding (region demarcated by a white line in Figure 1B) and the volatiles emanating from the OSP chamber. Mice made repetitive snout pokes into the orifices of the plate guarding the chambers to sense the stimuli. Such an experimental design facilitated the multimodal learning of pheromone locations with the stimuli. Thus, another parameter to measure the preference was the number of active attempts or snout pokes into the orifices of specific diameter to sense the volatiles. Decayed preference was observed over the course of 4 consecutive initial testing days. On day 2 of the initial testing phase, we observed an increase in the time spent towards OSP chamber indicating the emerging preference towards male urinary pheromones, which was reduced during the remaining two days (Figure 1C1: p > 0.05, F = 4.34 Repeated measures (RM) Two-way ANOVA). (Figure 1C1: RM Two-way ANOVA, Bonferroni’s multiple comparison test, p < 0.05 for day 2 and p > 0.05 for all other days). Similarly, on initial days, i.e. day 1 and day 2, female mice made higher number of active attempts to sample the urinary volatiles. The number of attempts, however, became similar to the exploration behavior on water side as the days progressed (Figure 1C2: p > 0.05, F = 4.1, RM Two-way ANOVA; Bonferroni’s multiple comparison test, p ≤ 0.01 for day 1 and day 2, p > 0.9 for day 3 and day 4). To test if this overall decline is due to variable responses shown towards both stimuli, we quantified and compared their attraction specifically towards the pheromones. Indeed, female mice spent lesser time (Figure S1A1: p < 0.01, F = 6.3, RM One-way ANOVA; Bonferroni’s multiple comparison test, p < 0.05 for day 2 v/s day 3 and day 2 v/s day 4) and made fewer active attempts particularly towards OSP chamber as the days progressed during the initial testing phase (Figure S1A2: p < 0.0001, F = 16.3, RM One-way ANOVA; Bonferroni’s multiple comparison test, p < 0.01 for day 1 v/s day 3, 4 and day 2 v/s day 3, 4).

In order to investigate if the association of non-volatiles and volatiles through olfactory and whisker systems can be strengthened over time, we further trained mice for a period of 15 days after their initial testing period was over. During this time, a mouse is allowed to enter only one of the zones and the door is then closed for 5 minutes ensuring the intended associative learning of pheromone locations with the volatiles and the specific hole size. This is then alternated for both sides leading to a total of 15 minutes being spent on each side. Indeed, this restriction to one particular zone allowed mice enough time to explore the hole size, sense the volatiles and the non-volatiles from soiled bedding. To investigate if such a multimodal association can be retained, we checked for pheromonal memory at day 15 and day 30 post training phase. Exposure to pheromonal stimuli was not allowed during the time period after the training. We observed an increased preference towards OSP chamber on day 15, as measured by the two parameters: time spent and number of active attempts (Figure 1D1, D2). This confirms that mice had intact memory at day 15 (Figure 1E1, p < 0.01 for time spent, Wilcoxon matched pairs signed rank test and Figure 1E2, p < 0.001, Paired two-tailed student’s t-test) after the multimodal training was carried out. However, we did not observe any preference when the 30^th^ day memory was analyzed (Figure 1E3: p > 0.05 for time spent and Figure 1E4: p > 0.1 for number of active attempts, Wilcoxon matched pairs signed rank test) after the training. As mice were not exposed to the stimuli for an extended period of 30 days, it possibly led to extinction of preferential memory for the attractive stimulus. This result is in line with other studies, which have shown that female mice can remember the scent location up to two weeks after a single exposure to the urine scent (14, 22). However, in their study, the reported memory was dependent on the spatial cue based learning.

To systematically investigate if whisker system is being utilized as a result of repeated snout insertions into the plate with orifices of specific diameter during the training phase, we carried out whisker sensory deprivation in another set of mice (Group 2, see Methods). In this experiment, whiskers were trimmed and an anesthetic gel was applied on the snout of the mouse during the training phase of 15 days so as to prevent the association of volatiles emanating from the stimulus kept inside the chambers guarded with plates of specific hole diameters. Application of anesthetic gel was done each day, 15 minutes before putting the mouse in the setup for the training period. Towards the end of this recovery period, we observed normal mobility with all animals tested. The mice were then put in the multimodal pheromonal learning set-up. Whiskers were not trimmed after the training period and all mice had re-growing whiskers on the day of memory investigation, Day 15 post training. The memory was significantly reduced in the whisker deprived group compared to whisker intact group indicating the involvement of whisker system in facilitating the multimodal learning and thereby the memory formation for pheromone locations (Figure 2A2: p < 0.0001, F = 18.69 Ordinary One-way ANOVA; Bonferroni’s multiple comparison test, p = 0.0008 for time spent and Figure 2B2: p < 0.0001, F = 15.27 Ordinary One-way ANOVA; Bonferroni’s multiple comparison test, p = 0.006 for number of active attempts). Indeed, we did not observe difference in the time spent in front of the OSP chamber v/s NS chamber (Figure S2A1: p > 0.1, Paired two-tailed student’s t-test) and the number of active attempts into the orifices on both the chambers (Figure S2A2: p > 0.1, Paired two-tailed student’s t-test) suggesting that preference for attractive, opposite sex pheromonal stimuli was impaired for this group. This corroborates to the importance of tactile cues in generating sexual-physiological changes, which has also been previously looked at. For instance, social touch along with the pheromones can lead to accelerated puberty in female mice suggesting the relevance of tactile cues (23).

Acquisition of memory of OSP location under whisker intact but not deprived conditions called for further interrogation. To study if the preference shown by female mice was specific to the opposite sex pheromones, we carried out multimodal training paradigm with “non-attractive”, same-sex pheromones using another set of female mice (Group 3, see methods). On testing the memory, we observed significantly lower response compared to the preference shown by Group 1 mice (Figure 2A2, B2: p < 0.0001 for both parameters, Ordinary One-way ANOVA, Bonferroni’s multiple comparison test). Further, there is a possibility that mice might adopt varying sampling strategies for the same stimuli under whisker intact and deprived conditions, which can modulate their response. We investigated sampling behavior using subsets of female mice utilized for multimodal pheromonal learning assay. We analyzed the sniffing frequencies shown by whisker intact and deprived mice towards the volatile pheromonal cues during brief pheromonal exposure epochs (Figure S3A). These mice were head-restrained and urine volatiles were delivered using a custom-built olfactometer. Their sniffing strategies did not vary under different conditions for the time duration we tested for (Figure S3B: p > 0.1 across all groups on four consecutive days, Ordinary One-way ANOVA). This confirms the fact that the reduced memory observed in Group 2 & 3 is not due to sampling differences, but because of the robustness and specificity of associations formed between different modalities.

A recent study indicates that an initial sexual experience can lead to gain in the population of excitatory Scnn1a expressing neurons in layer 4 of genital cortex in somatosensory areas in pubertal female mice (24). As a starting point towards depicting the role of subsystems involved in learning and memory of pheromonal locations in our paradigm, we explored the activation pattern of Arc protein on the last training day (TrD 15) and memory day 15 (MD 15) trained mice. Arc gene expression is dependent on the extent of neural plasticity achieved under certain behavioral and physiological conditions (25). Arc is activated by tasks involving active learning and exposure to novel stimuli (26). The spatio-temporal dynamics of Arc expression, compared to other immediate early genes suggests its role in reliably encoding memory traces (27, 28). We sacrificed the mice to obtain the brain within 0-15 minutes of their performance completion in the multimodal learning behavior on TrD 15 and MD 15. Arc immunoreactive cells were observed in the MOB both on TrD 15 and MD 15 of the whisker intact mice (Figure 3 A1-A4). An increasing trend in the Arc immunoreactive cell number on MD 15 indicates the role of MOB circuitry in governing learning and memory (Figure 3 A5, #, p = 0.072, Unpaired t-test, two-tailed). This could be due to the enhanced recruitment of OB adult-born granule cells (abGCs) in contributing towards the memory of pheromone locations. This needs to be further validated by looking at the Arc expression specifically in abGCs and further understanding their possible role in pheromonal memory retrieval.

We established that our pheromonal based learning and memory task can bring about activation of MOB circuitry. We further wished to assess if such an activation of OB circuitry can modulate other olfactory driven behaviors, we carried out a nonvolatile olfactory discrimination learning and memory tasks in another groups of female mice (Figure 3B). To this end, we carried out go/no-go odor discrimination paradigm (21) to investigate modulation of discrimination learning and memory of non-pheromonal volatiles in Whitten effect induced female mice (Figure 3B, C1-C3). CI v/s EU odor pair training was carried out for both the groups before Whitten effect was induced in experimental group to detect any possible bias due to the learning efficacy differences that might be existing between two groups (Figure 3D: CI v/s EU Ordinary Two-way ANOVA, Bonferroni’s multiple comparison test, p > 0.05 for all data points in learning curve). Induction and synchronization of estrous cycle, i.e. Whitten effect was achieved in a group of female mice by exposing them directly to male urine and bedding (Figure S4). Once the synchronization was achieved, these mice were trained on an odor-reward association task using the go/no-go odor discrimination paradigm. This was done to observe if MBU exposed mice can achieve quicker learning on an odor-reward association task as compared to mice who were just exposed to their female conspecifics in their cage. We did not find any differences in the learning pace for various odor pairs (Figure 3D: AA v/s EB: p > 0.1, F = 0.12; BZ v/s NN: p > 0.1, F = 1.43; HX v/s PN: p > 0.1, F = 0.19; Ordinary Twoway ANOVA, Bonferroni’s multiple comparison test, p > 0.05 across all data points for all learning curves). Individual day accuracies were also plotted for AA v/s EB odor pair to compare the day-wise accuracy level changes between the experimental and control groups for 4 continuous days (as the estrous cycle was of four days length in synchronized group) during the discrimination training days (Figure 3F: p > 0.1, F = 0.49, Ordinary Two-way ANOVA; Bonferroni’s Multiple comparison test, p > 0.9 across all data points). Discrimination accuracy levels were independent of the estrous stages mice were in. We also checked if the memory for one of the odor pairs (AA v/s EB) was enhanced when checked one month later after the discrimination training. MBU exposure leading to estrous synchronization did not lead to changes in non-pheromonal volatile discrimination behavior (Figure 3G: p > 0.1, Mann-Whitney test). These results indicate least influence from the estrous stages on the MOB dependent odor discrimination learning and memory at least with the paradigm we used. Our results from Figure 3 also demonstrate that female mice could very well be utilized for olfactory driven tasks and that probable sex-dependent changes in the behavioral readouts actually depend on the type of paradigm and the complexity of task employed.

Apart from MOB, we found that the Arc immunoreactive cells were also present in the somatosensory cortex and hippocampus of the MBU exposed whisker intact and deprived mice (Figure 4 E1 & F1). Such a specific activation profile of Arc indicates the involvement of olfactory and somatosensory neural circuits in pheromonal location learning & memory formation. Layer 4 of S1 cortex contain ‘barrels’ while the Layer 6 contain neurons forming ‘infrabarrels’ which are spatially aligned with barrels of layer 4. Distinctive neuronal types present in Layer 6 receive inputs from distinct thalamic sub-regions, thereby, capable of integrating information receiving from different pathways (29). We found out significantly higher number of Arc immunoreactive cells in the somatosensory cortical layers (Figure 4E1-E3) and the Dentate Gyrus (DG) of hippocampus (Figure 4F1-F3) in whisker intact group when compared to the whisker deprived group. Such a differential activation in intact v/s deprived groups correlates with the impaired acquisition and retrieval of pheromonal memory in the whisker deprived group (Figure 4E2, p < 0.0001; 4E3, p < 0.0001; 4F2, p = 0.005 and 4F3 p = 0.006; Unpaired t-test, two-tailed). In addition to this, we observed increased DG Arc activation in both the groups when probed on MD 15 as compared to TrD 15 (Figure S5C, p = 0.005 for whisker intact and p =0.007 for whisker deprived, Unpaired t-test, two-tailed). The role of hippocampus in encoding memory retrieval is thus, unscathed in deprived group, and it is primarily the lack of association that is happening between the olfactory and whisker subsystems which is not resulting in the formation of a robust, specific pheromonal memory (Figure S5A and S5B, p > 0.1 for whisker deprived group, Unpaired t-test, two-tailed).

**Figure 4.**
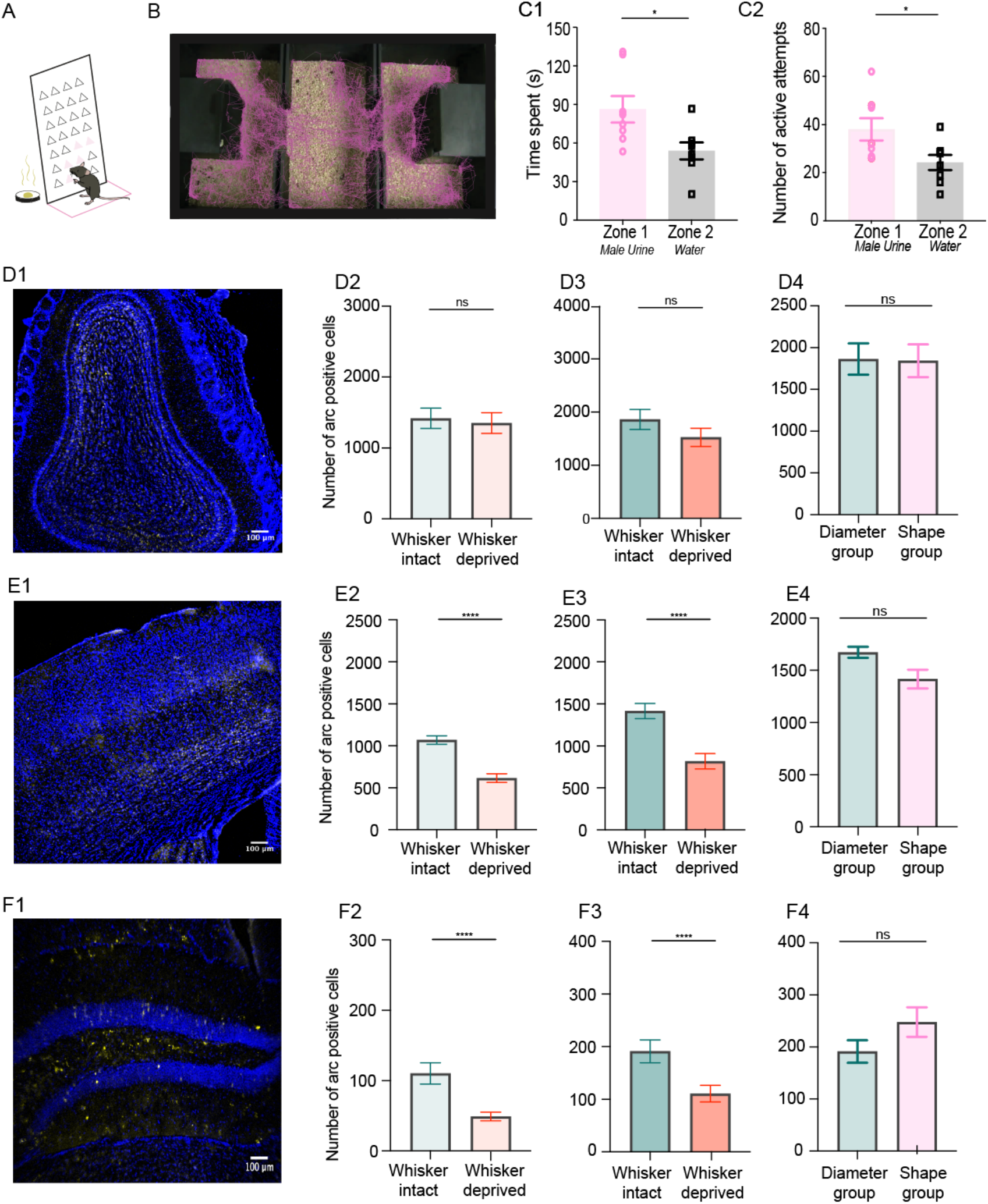
Long-term memory formation in a shape-based multimodal pheromonal learning paradigm is supported by enhanced Arc expression pattern. A. Schematic representing a female mouse sensing the male urine present inside the chamber through the triangle-shaped orifices on the plate guarding the chamber. The oppositely placed NS chamber is guarded with a plate with 10mm diameter circular orifices. B. Track recorded using EthoVision software by tracking the ‘nose point’ of the animal. Visibly more time spent near zone 1 compared to zone 2 on the day of memory testing (day 15^th^). C1. More time spent in zone 1 (Male Urine) by female mice on the 15^th^ day memory testing for shape based multimodal pheromonal learning task (p=0.013, Paired Student’s t-test, N = 8 mice). C2. Higher number of active attempts into the orifices (of either shape) on the plate guarding the male urine chamber checked on the 15^th^ day memory testing (p = 0.02, Wilcoxon test, N = 8 mice). D1. Arc immunoreactive cells in the MOB of the whisker intact female mouse exposed to MBU. D2. Similar number of Arc immunoreactive cells in the MOB of whisker intact and deprived mice on the last training day, i.e TrD15 (p = 0.749, Unpaired t-test, twotailed, N = 10-11 Region of interest (ROIs) from 2-3 mice, teal: Whisker intact and red: Whisker deprived). This is indicative of the similar sensory input and processing in these groups of mice as only somatosensation was blocked leaving the olfactory processing unaffected. D3. Similar number of Arc immunoreactive cells in the MOB of whisker intact and deprived mice on MD 15 (p = 0.204, Unpaired t-test, two-tailed, N = 10-11 ROIs from 3 mice). D4. Similar levels of Arc positive cells in the MOB of whisker intact female mice that carried out shape based multimodal pheromonal learning task when compared to mice that performed size/diameter based task (p > 0.05, Unpaired t-test, two tailed N = 8-11 ROIs from 2-3 mice). E1. Arc immunoreactive cells in the somatosensory cortical layers of the whisker intact female mouse exposed to MBU. E2, E3. Significantly higher number of Arc immunoreactive cells in the Somatosensory cortex (layer 4 and 6) in whisker intact group on TrD 15 (p < 0.0001, N = 12-18 ROIs from 2-3 mice) & MD 15 (p < 0.0001, Unpaired t-test, two-tailed, N = 18 ROIs from 3 mice). Differential Arc immunoreactivity is suggestive of the involvement of somatosensory cortical layers in the pheromonal location memory acquisition. E4. Similar levels of Arc positive cells in the somatosensory cortex of whisker intact female mice that carried out shape based multimodal pheromonal learning task when compared to mice that performed size/diameter based task (p > 0.05, Unpaired t-test, two tailed N = 7-8 ROIs from 2-3 mice). F1. Arc immunoreactive cells in the Hippocampus of the whisker intact female mouse exposed to MBU. F2, F3. Significantly higher number of Arc immunoreactive cells in the Hippocampus, a centre of learning & memory in case of whisker intact group on TrD 15 (p = 0.005, Unpaired t-test, two-tailed, N = 8-12 ROIs from 2-3 mice) and MD 15 (p = 0.006, Unpaired t-test, two-tailed, N = 12 ROIs from 2-3 mice). F4. Similar levels of Arc positive cells in the DG of whisker intact female mice that carried out shape based multimodal pheromonal learning task when compared to mice that performed size/diameter based task (p > 0.05, Unpaired t-test, two tailed N= 6-12 ROIs from 2-3 mice).

Both the groups received similar olfactory inputs during the testing and training session as only the whisker-mediated inputs were blocked in the deprived group. This suggests that similar olfactory processing would be occurring in both groups and clearly, we saw similar Arc activation in the OBs of these two groups of mice when checked on TrD 15 (Figure 3A5, p=0.7, Unpaired t-test, two-tailed).

To further substantiate the multimodal aspect of our paradigm, we also carried out shape-based pheromonal learning in another set of female mice (Group 4). To this end, we carried out training entailing mice to associate triangle v/s circular orifices with the stimuli present inside the chamber (OSP and NS) (Figure 4A, B). An increase in time spent and number of active attempts when checked on day 15^th^ after training indicated the presence of shape driven multimodal memory of pheromonal location (Figure 4C1 p=0.013, paired two-tailed student’s t-test, 4C2: p=0.02, Wilcoxon test). The coordinated action of sniffing pheromones and whisking through the orifices/triangles leading to memory acquisition & retention can point to the association between olfactory and whisker subsystems, which led us to investigating the Arc expression in this group of female mice as well. Similar number of Arc positive cells were found in the three brain regions (Figure D4, E4 & F4, p > 0.05 for all plots, Unpaired t-test, two-tailed) when compared to the mice who performed the size/diameter based multimodal pheromonal memory task (MD15). Our finding further corroborates to the involvement of associations between multiple sensory systems even while using a shape based multimodal pheromonal learning paradigm.

Overall, the multimodal aspect of the pheromone location learning leading to the preference memory formation of urinary pheromones suggests that pheromonal communication is not just a simple stimulus-response system. Even in humans the mediation of sexual attraction involves both tactile sensation and the attractive pheromonal detection (30). Further, we show that olfactory discrimination of non-pheromonal volatile odors is independent of the activation of pheromonal dependent MOB Arc activation that occurs upon male urine and bedding exposure. With respect to socio-sexual preference, female mice could exhibit enhancement of pheromonal location memory when more than one sensory system is mediating the associative learning. Our findings thus provide an evidence for formation of multimodal learning and memory in tasks pertinent to social and reproductive behaviors.

## Author Contributions

N.A. and M.P. carried out the study conceptualization and experimental design. M.P. and S.M. performed pheromonal location preference behavioral experiments with help from M.M.P. M.P. analysed the data acquired from pheromonal location preference behaviour. M.P. acquired and analyzed immunohistochemistry data. M.P., S.M. and U.D. performed and analyzed non-pheromonal volatile discrimination experiments. N.A. and M.P. wrote the manuscript with comments from S.M.

## Acknowledgements

We thank Thomas Kuner, N.K. Subhedar and Laboratory of Neural Circuits and Behaviour (LNCB) members for fruitful discussions. Part of the work was carried at the National Facility for Gene Function in Health and Disease (NFGFHD) at IISER Pune, supported by a grant from the Department of Biotechnology, Govt. of India (BT/INF/22/SP17358/2016). We thank staff of *National Facility for Gene Function in Health and Disease* (NFGFHD) and IISER Biology-Leica microscopy facility for the technical support. This work was supported by the DBT/Wellcome Trust India Alliance intermediate grant (IA/I/14/1/501306 to N.A.), IISER-Pune Fellowship (M.P.) and CSIR Fellowship (S.M.).

## Supplementary Figures

**Supplementary Figure S1.**
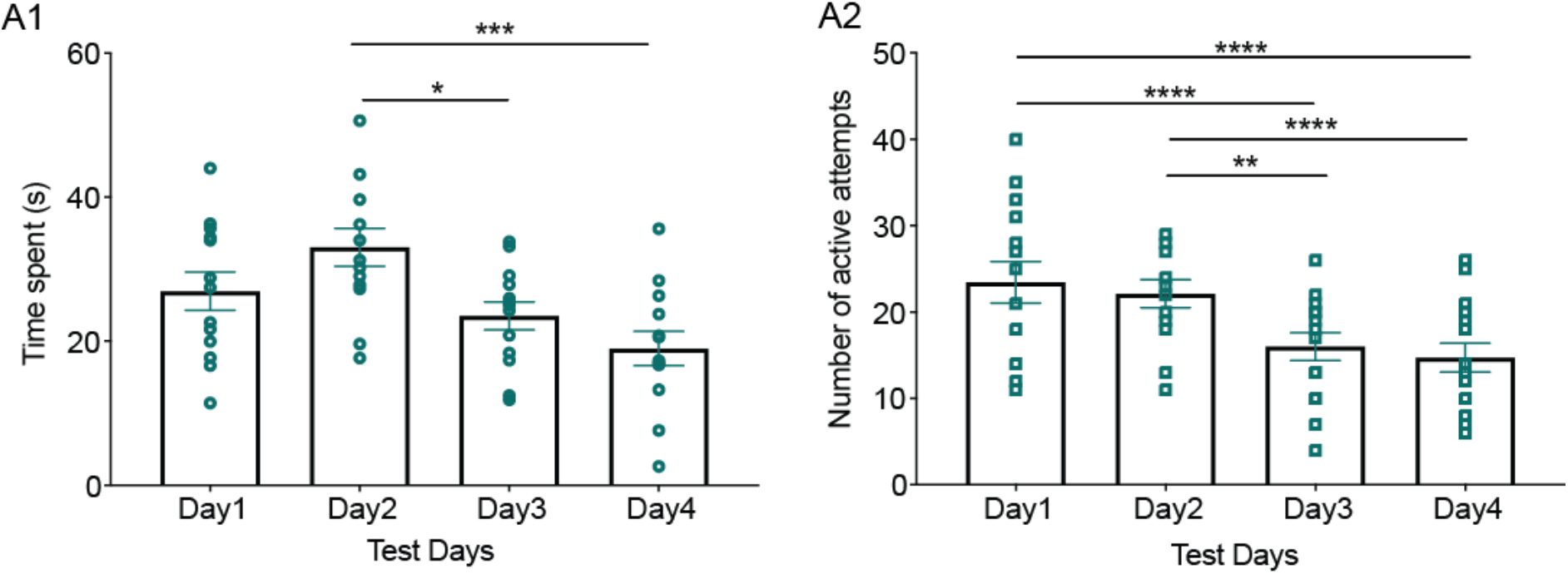
Decayed preference for pheromone locations in whisker-intact female mice exposed to a choice of male urine and neutral stimulus. A1. Time spent near OSP chamber containing male urine and soiled bedding by whisker intact female mice. Decayed preference towards this chamber was observed during the initial testing phase (p < 0.01, F = 6.3, RM One-way ANOVA; Bonferroni’s multiple comparison test, p = 0.04 for day 2 v/s day 3 and p = 0.001 for day 2 v/s day 4; p > 0.1 for all other comparisons, N = 13 mice). A2. Number of active attempts towards OSP chamber (male urine) during the initial 4 days of testing phase. Mice showed lower number of nose pokes on day 3 and 4 (p < 0.0001, F = 16.3, RM One-way ANOVA; Bonferroni’s multiple comparison test, p < 0.0001 for day 1 v/s day 3, 4 and p < 0.001 day 2 v/s day 3, 4; p > 0.9 for all other comparisons, N = 15 mice).

**Supplementary Figure S2.**
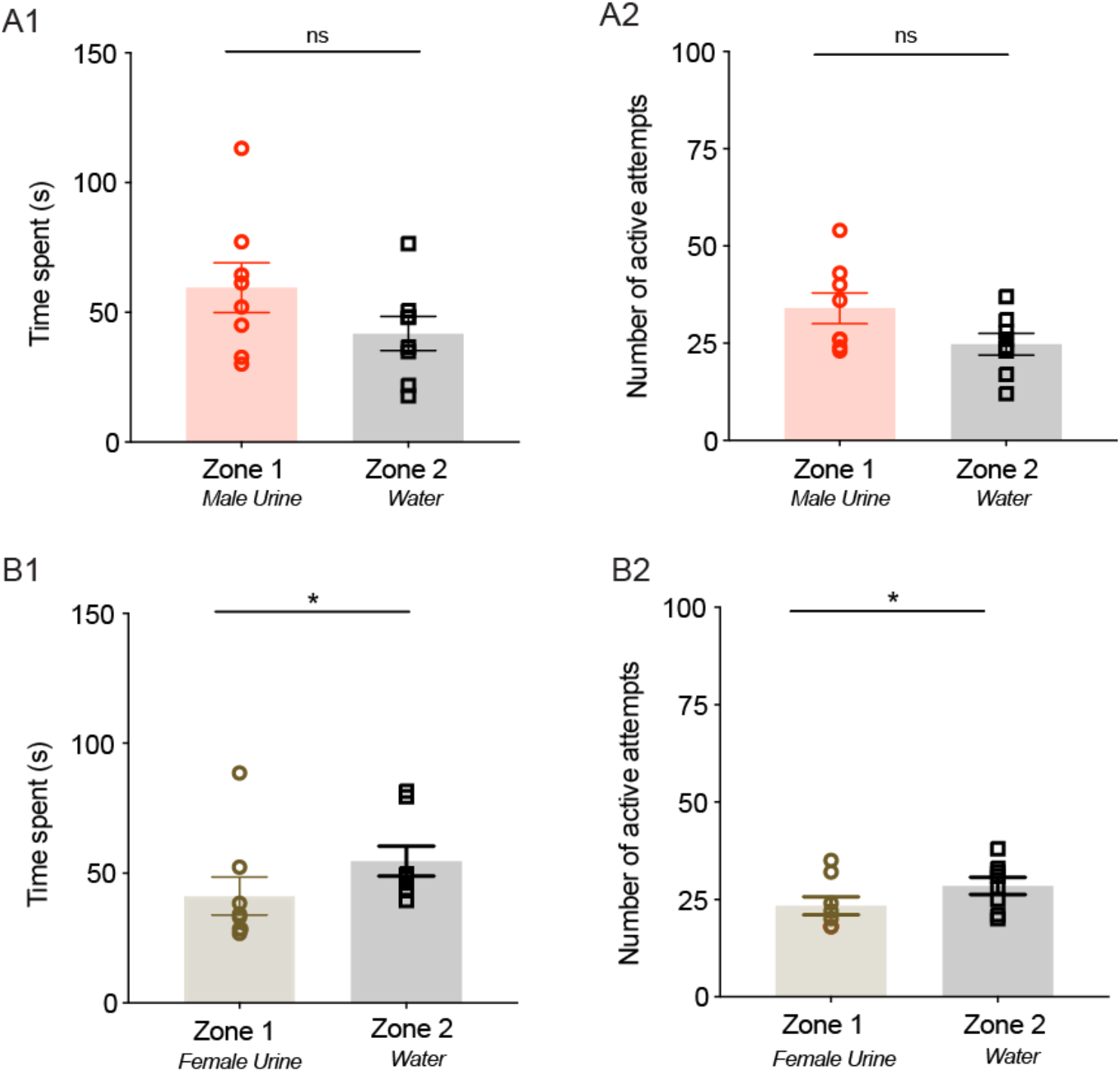
Similar sampling behavior shown towards pheromonal and neutral stimuli. A1, A2. No preference for OSP as indicated by similar time spent and number of active attempts for whisker deprived female mice on day 15^th^ memory phase. The mice were trained with male urine and bedding v/s water. (p > 0.1 for time spent and for number of active attempts, Paired two-tailed student’s t-test, N =8 mice) B1, B2. Decreased preference for female urine as indicated by increased time spent and number of active attempts on NS chamber for whisker intact female mice on day 15^th^ memory phase. The mice were trained with female urine and bedding v/s water. (p = 0.039 for time spent and number of active attempts, Wilcoxon matched pairs signed rank test, N = 8 mice)

**Supplementary Figure S3.**
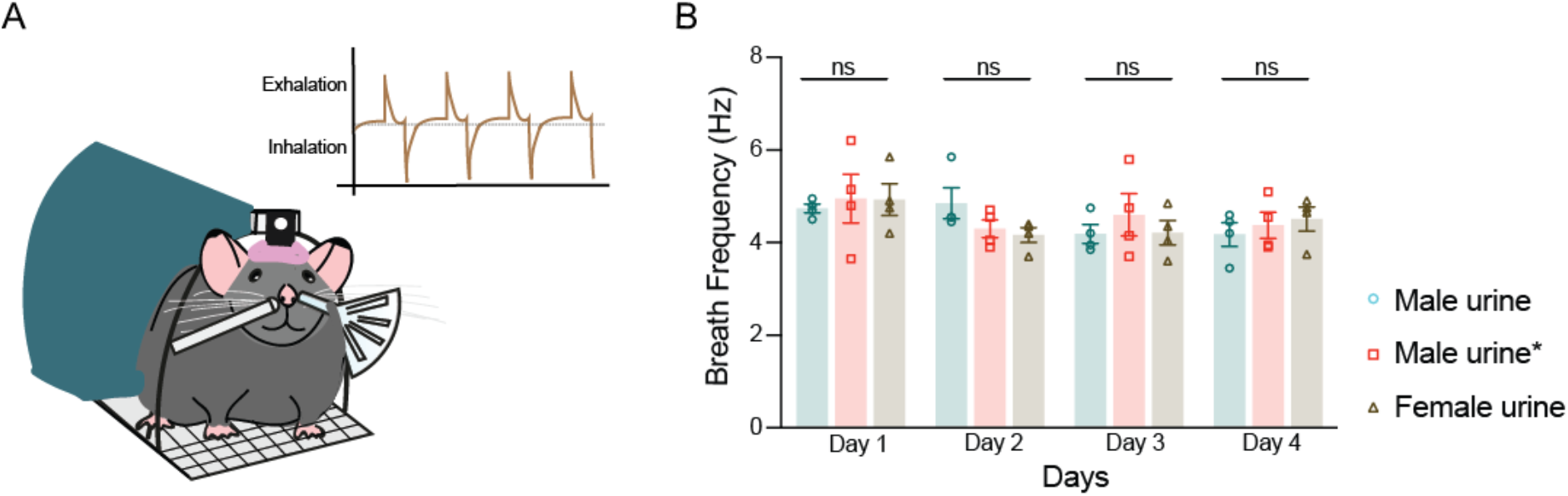
Similar sniffing frequency for sampling pheromones under whisker intact and deprived conditions. A. Schematic of a female mouse head-post implanted on the head-restrained setup. Breathing is recorded using air-pressure sensor placed near one nostril (on left nostril in depiction) and delivering the stimulus (male urine or female urine volatiles) pseudorandomized with filtered air presentations from a nozzle (right side). Inset: Representative Inhalation/Exhalation pattern of a mouse breathing during stimulus delivery. B. Breath frequency in three groups of female mice (blue: whisker intact sensing male urine volatiles, red: whisker deprived, anesthetic applied sensing male urine volatiles and yellow: whisker intact sensing female urine volatiles) recorded across four consecutive days (day 1: p = 0.9, F = 0.1; day 2: p > 0.1, F = 2.2; day 3: p = 0.6, F = 0.58; day 4: p = 0.7, F = 0.40; Ordinary One-way ANOVA, Bonferroni’s multiple comparison test, p > 0.1, N = 4 mice for all groups) (Male urine*: whisker deprived group/Group 2).

**Supplementary figure S4.**
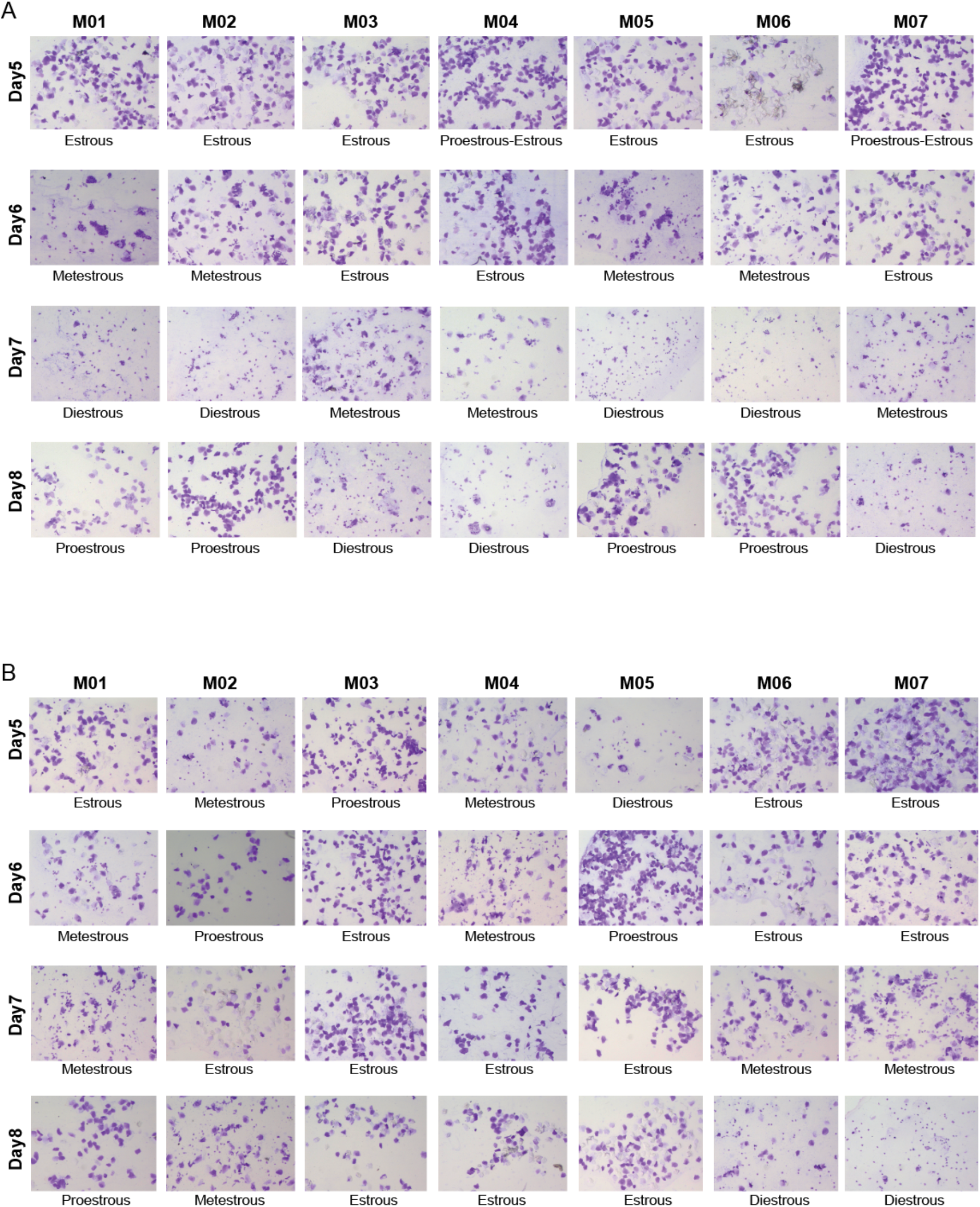
Whitten effect induction and synchronization in female mice by male pheromonal exposure. A. Whitten effect, i.e. induction and synchronization of estrous cycle continued to occur in experimental female mice during the duration of AA v/s EB odor pair training. Subset of mice whose vaginal smears were taken on a daily basis during the training, four consecutive days of induction and synchronization is observed in case of experimental mice (each stage lasts for a day, making it a proper 4 day cycle and follows an expected sequence in cycle from proestrous -> estrous -> metestrous -> diestrous). B. Control group which were not exposed to male urine and soiled bedding exhibited prolonged estrous cycle with one stage lasting > 1 day and not occurring the proper sequence of the cycle, i.e. deviating from proestrous -> estrous -> metestrous -> diestrous sequence.

**Supplementary Figure S5.**
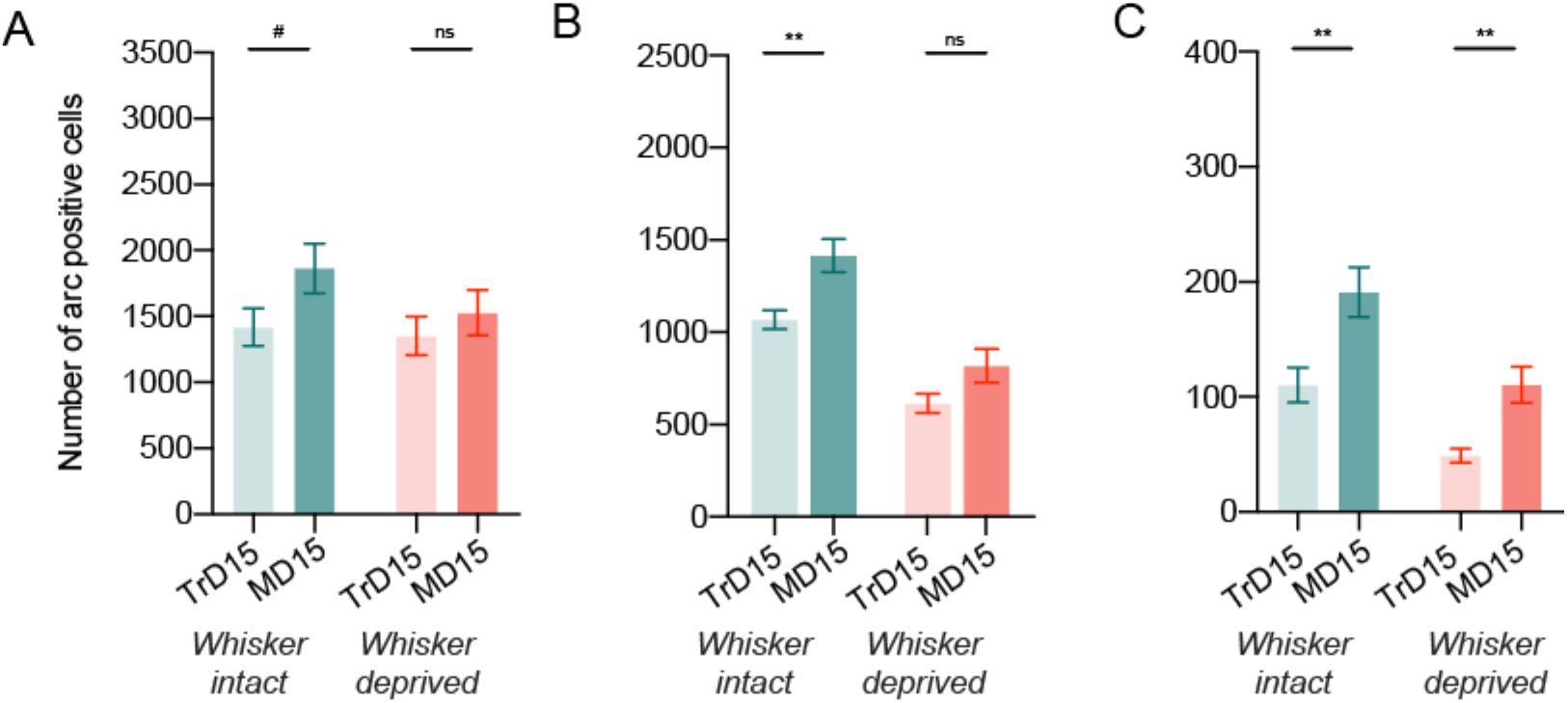
Trend of Arc activation during training and memory in the multimodal pheromonal learning paradigm. A. A slight yet non-significant increase in proportion of Arc immunoreactive cells in whisker intact group from TrD 15 to MD15 (#, p = 0.072, Unpaired t-test, twotailed, N = 10-11 ROIs from 2-3 mice). Similar proportion of Arc immunoreactive cells in whisker deprived group was found across TrD 15 and MD 15 (p =0.44, Unpaired t-test, two-tailed, N = 11 ROIs from 2-3 mice). B. Increase in Arc-positive cells in the somatosensory cortex of the whisker intact group on the Memory day 15^th^ when compared to last day of training (p = 0.0019, Unpaired t-test, two-tailed, N = 18 ROIs from 3 mice) while no change is seen in whisker deprived group (p = 0.107, Unpaired t-test, two-tailed, N = 12-18 ROIs from 2-3 mice) which evidences towards the involvement of whisker system in acquiring and retaining the multimodal pheromonal memory. C. Increase in Arc positive cells of Dentate Gyrus, Hippocampus in both the groups when compared from TrD 15 to MD 15 (p = 0.005 for whisker intact, N = 12 ROIs from 2-3 mice and p =0.007 for whisker deprived, N = 8-12 ROIs from 2-3 mice, Unpaired t-test, two-tailed). Such an increase indicates that the role of hippocampal circuits in memory retrieval is maintained normally even in whisker deprived group. Thus, it could primarily be the decreased activation of somatosensory cortex, which resulted in absence of memory acquisition in the deprived group.

